# The DNA methylation landscape of five pediatric-tumor types

**DOI:** 10.1101/2021.08.02.454814

**Authors:** Alyssa C. Parker, Badi I. Quinteros, Stephen R. Piccolo

**Affiliations:** Department of Biology, Brigham Young University, Provo, UT, USA

**Author notes:** Please address correspondence to S.R.P. at.

## Abstract

Fewer DNA mutations have been identified in pediatric tumors than adult tumors, suggesting that alternative tumorigenic mechanisms, including aberrant DNA methylation, may play a prominent role in pediatric tumors. Methylation is an epigenetic process of regulating gene expression in which methyl groups are attached to DNA molecules, often in promoter regions. In Wilms tumors and acute myeloid leukemias, increased levels of epigenetic silencing have been associated with worse patient outcomes. However, to date, researchers have studied methylation primarily in adult tumors and for specific genes but not on a pan-pediatric cancer scale. We addressed these gaps first by aggregating methylation data from 309 noncancerous samples and establishing baseline expectations for each gene. Even though these samples represent diverse tissue types and population ancestral groups, methylation levels were highly consistent for most genes. Second, we compared tumor methylation levels against these baseline values for five pediatric cancer types—Wilms tumors, clear cell sarcomas of the kidney, rhabdoid tumors, neuroblastomas, and osteosarcomas. Hypermethylation was more common than hypomethylation—as many as 11.8% of genes were hypermethylated in a given tumor, compared to a maximum of 4.8% for hypomethylated genes. For each cancer type, genes with the highest variance exhibited consistently divergent methylation patterns for distinct patient subsets. We evaluated whether genomic and epigenomic abnormalities contribute to pediatric tumorigenesis in a mutually exclusive manner but did not find evidence of this phenomenon. Furthermore, even though oncogenes are commonly upregulated in tumors, and tumor-suppressor genes are commonly downregulated in tumors, we did not find statistical evidence that methylation drives such patterns on a broad scale in pediatric tumors.

## Introduction

Pediatric tumors are the leading cause of disease-related death for children in developed countries^1^, and those who survive pediatric cancer often experience adverse health challenges later in life^2^. Many mutations and structural variants have been associated with adult forms of cancer^3^; however, significantly fewer genomic abnormalities have been identified in pediatric tumors^4^. The mutation rate in pediatric tumors is 14 times less than the mutation rate in adult tumors, implying that different mechanisms may be involved in pediatric-cancer development than in adult cancers^5^. Of the mutations that have been identified in pediatric tumors, many are associated with epigenetic regulation^1^. In many pediatric tumors, molecular profiling does not identify genomic abnormalities but does show abnormal DNA methylation patterns^6,7^, suggesting the DNA methylation may play a critical role in tumorigenesis in these cases.

Gene-expression levels in cancer cells often vary from those in normal cells^8^. Such abnormalities alter cellular environments and manipulate cellular processes, leading to increased survival, rapid proliferation, and metastasis^9^. Oncogenes are one type of genes that contribute to the development of these abnormal features. These genes are often expressed at higher levels in cancer cells than in normal cells^10^. Another type of genes known as tumor suppressor genes counteracts cellular changes that lead to cancer. These genes are often expressed at lower levels in tumor cells than in normal cells^11^, potentially leading to rapid cellular proliferation. Methylation of the promoter region is often negatively correlated with gene expression levels, suggesting that DNA methylation plays a role in regulating gene expression^12,13^. However, relatively little is known about global methylation patterns in pediatric tumors, the interplay between methylation events and mutations in pediatric tumors, or how these observations may differ between known cancer genes (oncogenes and tumor suppressors) and other genes. Prior studies have focused on cancer cell lines, a single tumor type at a time, or adult cancers^14–16^.

Computational models have been developed to identify differentially expressed genes across large sets of methylation data^15,17^. Applications of these methods have found several genes with highly variant methylation in adult tumors^17^. Many of the genes that exhibited highly variant methylation in tumors were not previously associated with cancer, and genomic aberrations in these genes were not characteristic of tumors. A small-scale study of Wilms tumors (a common pediatric cancer) showed similar results^17^. It has been shown that several distinct types of cancer, including endometrioid adenocarcinomas and glioblastomas, share many differentially methylated regions, suggesting that these epigenetic markers may be a universal feature of cancer^18^.

While these findings hint that aberrant methylation patterns may also be characteristic of pediatric cancer, most pediatric cancers have not been analyzed for DNA methylation patterns. One study investigated genomic, transcriptomic, and epigenomic patterns in acute myeloid leukemia and identified dozens of hypermethylated genes and age-specific patterns^19^. Additionally, structural variations were found to be more common than single nucleotide polymorphisms. A separate analysis of acute lymphoblastic leukemia identified a number of genes on chromosome 3, including PPP2R3A, THRB and FBLN2, that were frequently hypermethylated^16^.

Little work has been done to specifically investigate aberrant methylation of oncogenes and tumor suppressor genes in cancer. One study about endometrial cancer identified seven oncogenes that were hypomethylated and upregulated and twelve tumor suppressor genes that were hypermethylated and downregulated^14^, suggesting that changes in DNA methylation impact gene expression and may target oncogenes and tumor suppressor genes.

We address these gaps by analyzing methylation data for five types of pediatric cancers: Wilms tumors, clear cell sarcomas of the kidney, rhabdoid tumors, neuroblastomas, and osteosarcomas. We compare these cancer types against each other. Furthermore, as a baseline reference, we compare the tumor data against methylation levels for fetuses and children representing normal conditions for diverse cell types and human populations. Because methylation events tend to be gene specific^20^, we evaluate gene-level patterns. But we also consider global methylation patterns. For many of the tumors, we identify genomic alterations—single-nucleotide variants, small indels, and structural variants—in the tumors and evaluate whether these somatic mutations exhibited gene-specific mutual exclusivity with hypo- or hypermethylation events. In addition, we evaluate the consistency of these findings for oncogenes and tumor suppressor genes.

## Results

Our goal was to evaluate gene-level DNA methylation patterns for pediatric-tumor cells and normal cells. We used publicly available data to characterize five types of pediatric cancers as well as baseline methylation levels for normal cells. First, we evaluated the consistency of methylation levels representing non-cancerous states in fetuses and children. Second, we compared tumor methylation levels against the normal values and identified genes and tumor samples that exhibited patterns of hypomethylation or hypermethylation. Next, under the assumption that tumors with aberrant methylation would be subject to evolutionary constraints that are redundant with those resulting from somatic mutations, we evaluated whether these two event types were mutually exclusive in a given tumor. Finally, we examined these patterns within known oncogenes and tumor-suppressor genes.

### Consistency of methylation levels in normal samples and in pediatric tumors

To aid in understanding how methylation levels change in cancer cells, we first characterized baseline methylation levels for individual genes. We obtained Illumina Infinium 450K data for four normal datasets. We used data from diverse cell types and human populations with the goal of identifying methylation patterns that broadly represent baseline methylation states for healthy children. The datasets were from chorionic villi, kidney, spinal cord, brain, muscle, nasal epithelial, and blood cells and included data for a total of 309 patients of North American (n = 94), African American (n = 36), and Australian ancestry (n = 179).

Because DNA methylation contributes to regulating gene expression, we expected most genes to exhibit a consistent baseline methylation range. We anticipated that many genes (such as tumor suppressor genes) would have consistently low methylation levels because those genes must be consistently expressed to properly regulate cellular division, growth, and other proliferation activities. We anticipated that other genes (such as oncogenes) would have consistently high levels of methylation because proper cellular functioning requires that these genes remain relatively inactive. We expected to see some variance across samples because we included data from multiple cell types and because methylation levels change as cells respond to internal and external cues. We also anticipated that some genes would deviate from these patterns, perhaps in part because they are regulated by mechanisms other than DNA methylation. Based on a preliminary inspection of the data, we identified thresholds for categorizing genes based on the magnitude and variance of methylation levels. We considered genes with a median value less than 0.2 across all samples in a given dataset to be methylated at “low” levels, genes with a median greater than 0.6 to be methylated at “high” methylation levels and the remaining genes as having “medium” methylation levels. We categorized genes with a coefficient of variation (CV) less than 0.5 in a given dataset as having “low” variance and the remaining genes as having “high” variance. In the largest normal dataset (GSE89278), most genes exhibited low (58.9%) or medium methylation levels (25.6%). Relatively few genes exhibited high methylation (15.5%), meaning that under normal conditions, most genes appear not to be subject to strong expression constraints as a result of DNA methylation. Nearly all genes (95.1%) exhibited low variance. The other normal datasets reflected similar patterns.

By combining these two ways of categorizing the genes (Additional Data File S1), we found that the most common combination across all datasets was low methylation / low variance (56.9-57.2% per dataset). For GSE89278, the following numbers of genes fell into each category: low methylation / low variance: 12,874; low methylation / high variance: 534; medium methylation / low variance: 3,898; medium methylation / high variance: 14; high methylation / low variance: 5,246; high methylation / high variance: 0. These results were similar for the other normal datasets. To explore the consistency of these patterns across the normal datasets, we classified each gene into the following consistency levels: consistent (same category in all four datasets), semiconsistent (same category in three datasets), and inconsistent (same category in two or fewer datasets). Of the 22,253 genes that we profiled, 20,089 were consistent, 1,512 were semiconsistent, and 652 were inconsistent (Figure 1). Thus even though the normal datasets differed based on cell type and population ancestral groups, gene-level methylation levels were largely consistent.

**Figure 1:**
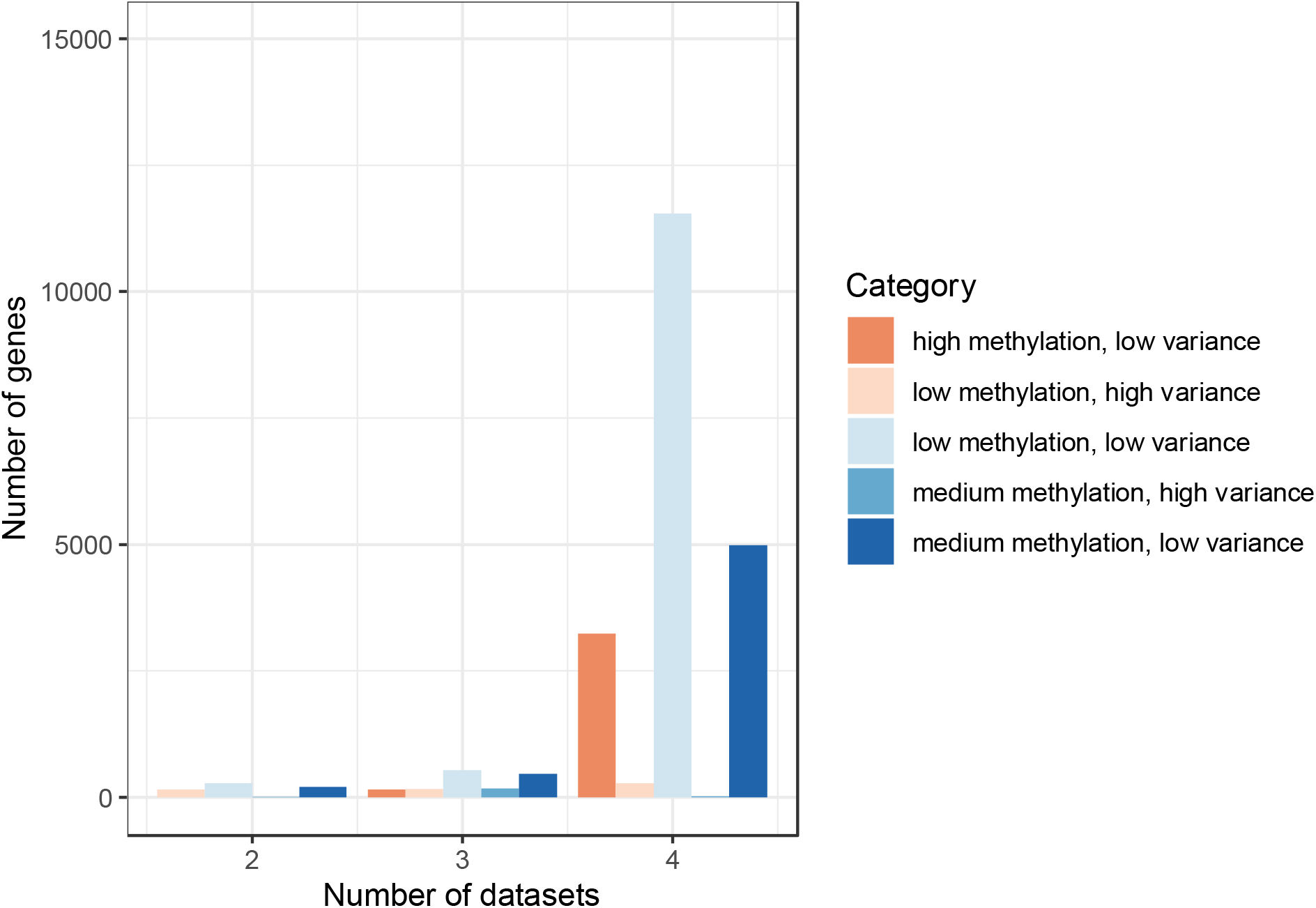
Consistency of DNA methylation levels and variances in normal cells. We assigned each gene to a category that indicated whether it was methylated at low, medium, or high levels and whether it had low or high variance across samples in a given dataset. For each gene, we counted the maximum number of datasets for which the level / variance category was consistent. For most genes, the category was consistent across all four datasets.

The gene with the smallest variance across the normal datasets was BTG3 (variance = 2.09E-5). In each of the normal datasets, BTG3 exhibited low methylation, suggesting that a biological constraint suppresses methylation of this gene under normal circumstances. The process that normally keeps BTG3 methylation low may be disrupted in tumors. Indeed, hypermethylation of the promoter region of BTG3 has previously been associated with several types of cancer, including breast^21^, prostate^22^, and renal^23^. The gene with the largest variance across the normal datasets was DOCK11. Information about this gene is sparse in the literature. Its role in cancer^24^ and hypercholesterolemia^25^ has been discussed. Its high variance in normal cells suggests that consistent expression of DOCK11 is inessential to normal cellular function and/or that processes other than promoter methylation regulate its expression.

### Tumor methylation relative to normal levels

Under the hypothesis that tumorigenesis alters local and global methylation patterns, we evaluated the extent to which methylation levels and variances differed for a given gene between normal and tumor conditions. For each gene, we compared the most common median / variance category across the four normal datasets against the most common category across the five cancer datasets. Typically genes stayed under the same category. For example, of the 12,368 genes that were categorized as low methylation / low variance for the normal datasets, only 549 (4.4%) changed categories: either to low methylation / high variance (n = 462) or to medium methylation / low variance (n = 87) (Table 1). The 227 genes in the medium methylation / high variance category were most likely to change categories, with 137 genes (60.0%) changing to low methylation / high variance and 19 genes (8.4%) changing to medium methylation / low variance. For the 512 genes that changed median methylation levels, approximately half (n = 259, 50.6%) increased (from low to medium or medium to high). For the 562 genes that changed variance categories, 519 (92.3%) increased from low to high variance. These increased variances suggest that the factors that normally keep methylation levels stable under normal conditions often become dysregulated in tumors. Altered expression of these genes may play a role in tumor development or may lead to downstream effects that affect tumor development. For example, if methylation levels of an oncogene are decreased, higher expression of the gene may result, leading to increased cellular growth, proliferation, or survival^9^. On the other hand, increased methylation levels of a tumor suppressor gene could cause lower gene expression and prevent it from performing regulatory functions^13^.

**Table 1:**
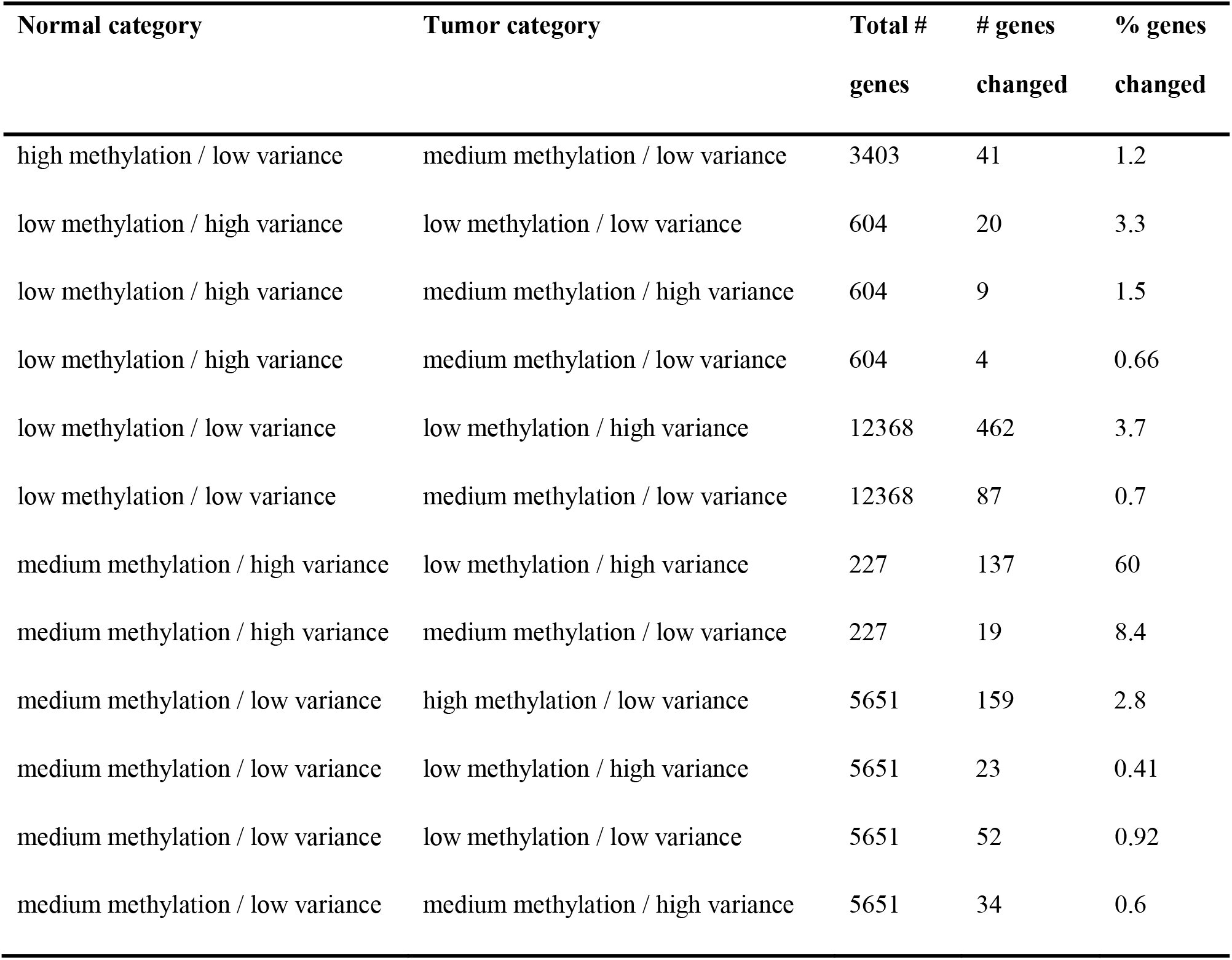
Summary of changes in methylation level/variance categories between normal and cancer datasets. We assigned each gene to a category that indicated whether it was methylated at low, medium, or high levels and whether it had low or high variance across samples in a given dataset. The table shows the total number of genes in each category for the normal datasets and the number (and percentage) of genes that changed from one category to another in the tumor datasets.

| Normal category | Tumor category | Total #<br>genes | # genes<br>changed | % genes<br>changed |
| --- | --- | --- | --- | --- |
| high methylation / low variance | medium methylation / low variance | 3403 | 41 | 1.2 |
| low methylation / high variance | low methylation / low variance | 604 | 20 | 3.3 |
| low methylation / high variance | medium methylation / high variance | 604 | 9 | 1.5 |
| low methylation / high variance | medium methylation / low variance | 604 | 4 | 0.66 |
| low methylation / low variance | low methylation / high variance | 12368 | 462 | 3.7 |
| low methylation / low variance | medium methylation / low variance | 12368 | 87 | 0.7 |
| medium methylation / high variance | low methylation / high variance | 227 | 137 | 60 |
| medium methylation / high variance | medium methylation / low variance | 227 | 19 | 8.4 |
| medium methylation / low variance | high methylation / low variance | 5651 | 159 | 2.8 |
| medium methylation / low variance | low methylation / high variance | 5651 | 23 | 0.41 |
| medium methylation / low variance | low methylation / low variance | 5651 | 52 | 0.92 |
| medium methylation / low variance | medium methylation / high variance | 5651 | 34 | 0.6 |

To identify genes that may influence pediatric tumorigenesis, we compared tumor methylation levels for each gene against the respective normal values on a per-cancer basis. We used a two-sided, Mann-Whitney U test and adjusted for multiple tests using the Benjamini-Hochberg False Discovery Rate (FDR)^26^. We considered genes with an FDR < 0.05 and an absolute methylation change > 0.02 to be statistically significant. Out of the 22,253 genes we analyzed, 37 exhibited differential methylation for at least one cancer type. Of these genes, 19 had decreased methylation in tumors, and 18 had increased methylation, including 15 genes that were differentially methylated for Wilms tumors, 0 for clear cell sarcomas, 6 for rhabdoid tumors, 17 for neuroblastomas, and 4 for osteosarcomas; 4 genes were statistically significant for 2 or more cancer types (Table S1; Figure 2). We note that the number of significant genes was smaller for cancer types with relatively small sample sizes.

**Figure 2:**
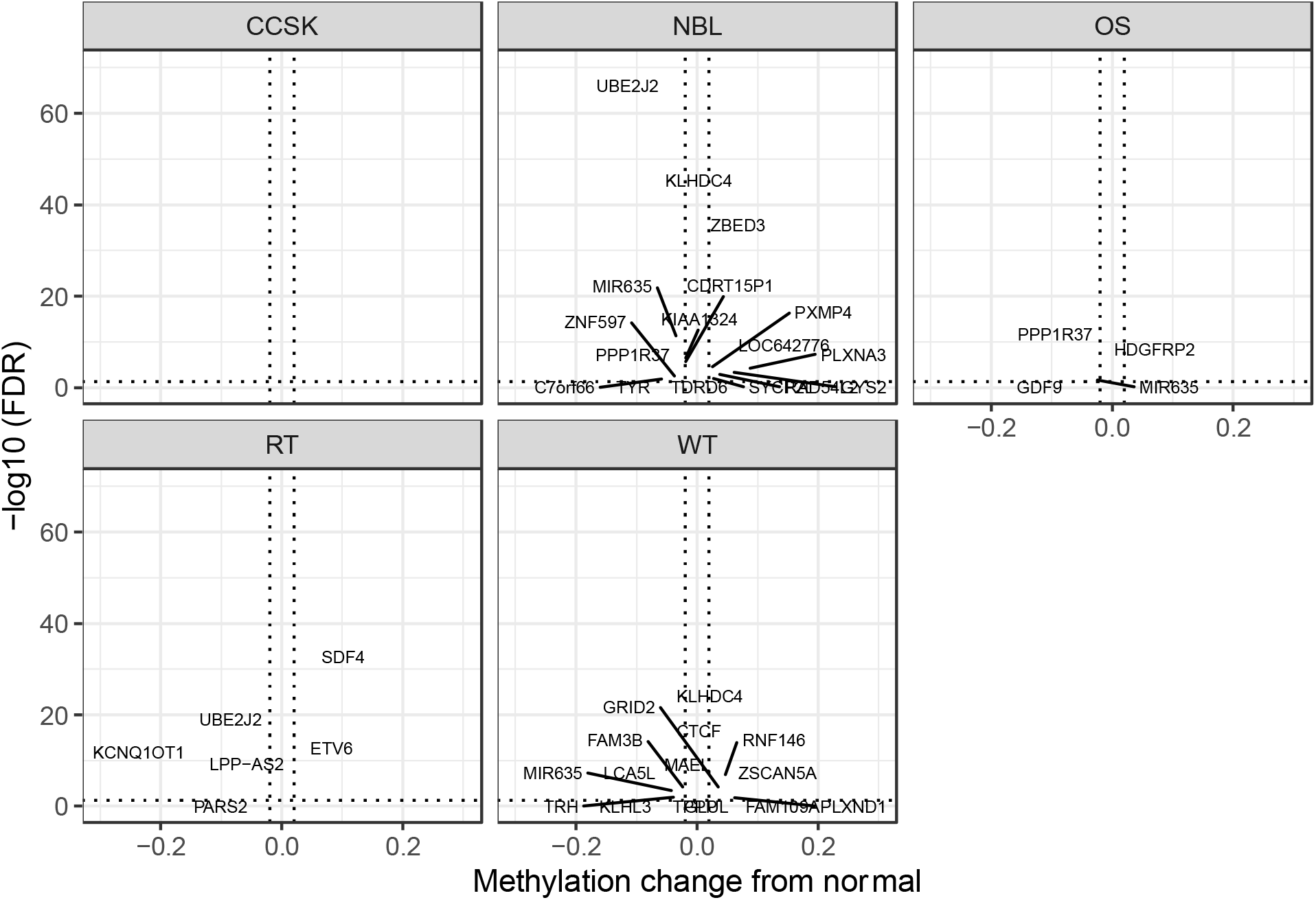
Volcano plots showing differentially methylated genes for each tumor type. For each tumor type, we compared methylation levels at the gene level between tumors and the normal samples. Genes showing significantly different methylation levels between tumor and normal conditions are highlighted.

We performed a pathway analysis for the significant genes from each cancer type using reactome.org^27,28^. For Wilms tumors, rhabdoid tumors, and osteosarcomas, pathways associated with ubiquitination, transcriptional regulation, and growth signaling were significant; for neuroblastomas, growth-signaling, integrin-signaling, and blood-biosynthesis pathways were among the most significantAdditional Data Files S2-S5. These functions have plausible connections to tumor biology because they are central to cellular function in general,^29,30^ but it is difficult to make precise inferences due to the relatively small numbers of significant genes.

To characterize differences in methylation across the five cancer types, independent of normal methylation levels, we performed a one-way ANOVA test for each gene and adjusted the p-values using the FDR correction. Methylation levels of most genes were consistent across the 5 cancer types, including for genes that we had identified as being differentially methylated in tumors compared to normal cells. Methylation levels for 19 genes differed significantly across the cancer types (FDR < 0.05; Table S2; Figure 3). But in most cases, the mean differences were small. For example, the largest absolute difference in mean methylation between tumor types for any of the significant genes was an increase of 0.24 in CCSK compared to RT (KCNQ1OT1 gene). Of the 19 genes, 10 fell into the medium methylation / low variance category, a higher proportion (0.526) than the overall proportion of genes in this category (0.175).

**Figure 3:**
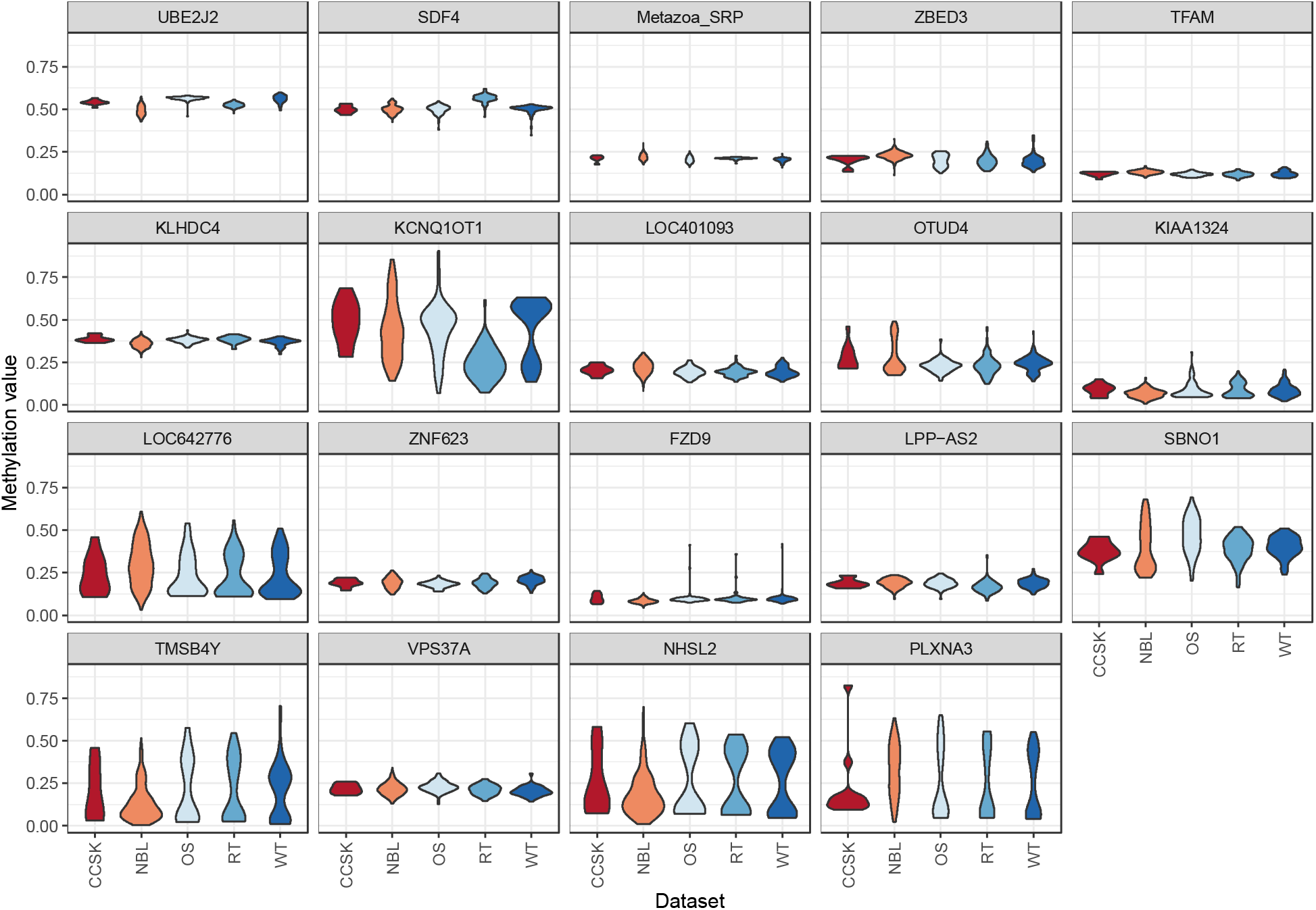
Methylation levels for genes that differed significantly across the tumor types. One-way ANOVA tests applied to tumor methylation levels identified 19 genes for which the means differed significantly across the tumor types. These violin plots show the range and density of the methylation values for these genes across the tumor types.

Next we focused on the 20 genes for which methylation levels varied most across all tumor types and plotted the methylation values, relative to normal levels, for each patient (Figure 4). Some tumors exhibited relatively high methylation levels for nearly all of these genes, whereas other tumors exhibited relatively low methylation levels for the same genes. This was especially true for Wilms tumors, rhabdoid tumors, and osteosarcomas. For example, the average methylation difference (relative to normals) for these 20 genes was 0.13 for 58 (44.3%) of the Wilms tumors but -0.11 for the remaining tumors. For the osteosarcomas and rhabdoid tumors, we observed a similar pattern in which subsets of 49 (57.0%) and 35 (51.5%) tumors, respectively, showed markedly higher methylation levels than the remaining tumors.

**Figure 4:**
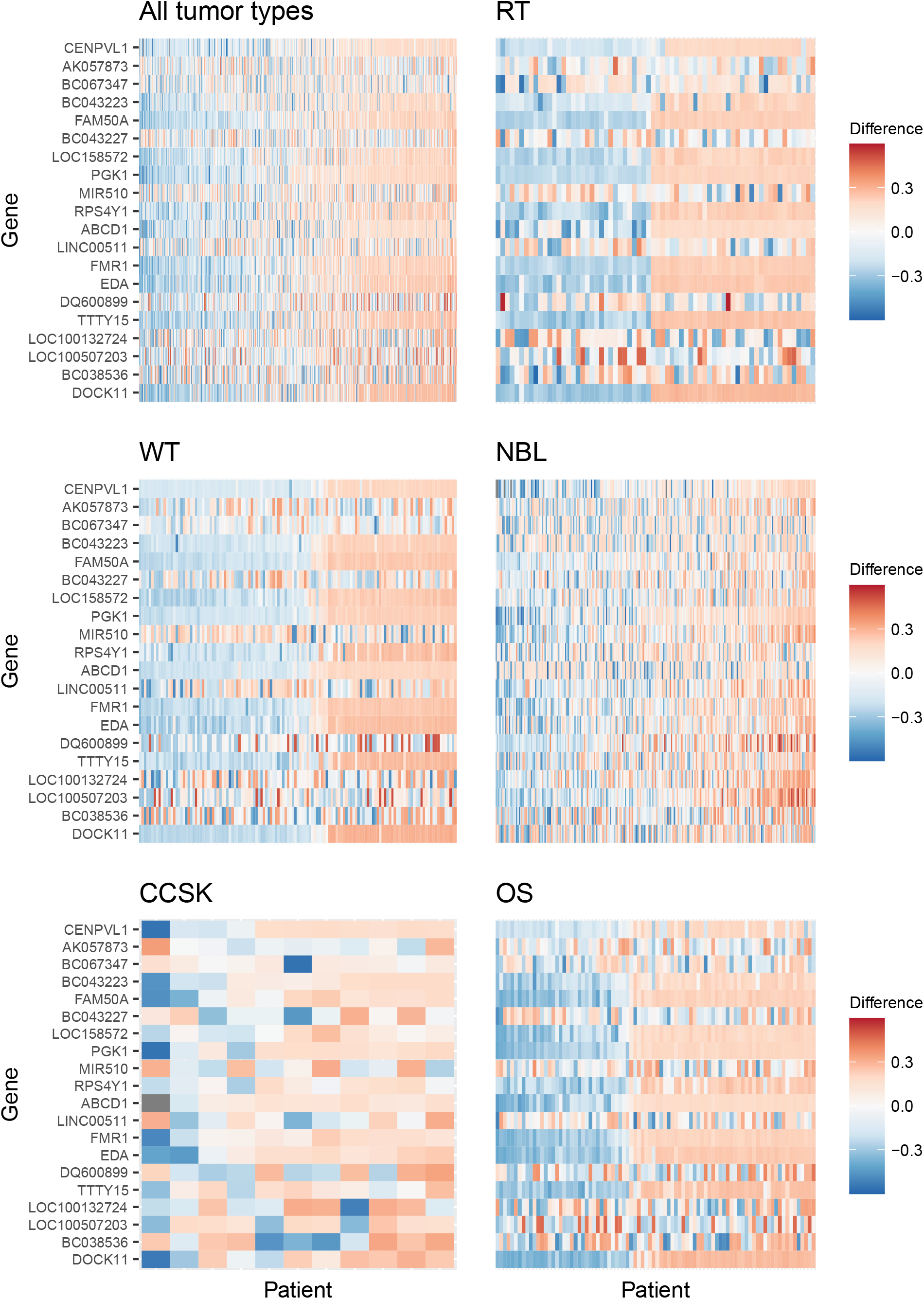
Gene-level DNA methylation changes for high-variance genes. Rows in these heatmaps indicate methylation levels, relative to the normal data, for the 20 genes with the largest variance across the tumor types. Columns represent individual tumors. Approximately half of all tumors exhibited consistently lower methylation levels than the remaining tumors.

These patterns of consistently divergent methylation might be useful for identifying tumor subtypes in a precision-medicine context^31^ and may be driven by factors such as the presence of somatic mutations and structural alterations that affect global methylation in characteristic ways.

To investigate global methylation patterns, we calculated gene-level z-scores for each tumor using the normal data as a reference. In cases where a tumor’s methylation value was more than three standard deviations higher than the mean normal value for a particular gene, we classified that gene as being hypermethylated in that tumor. In cases where a tumor’s methylation value was more than three standard deviations lower than the mean normal value for a particular gene, we classified that gene as being hypomethylated in that tumor. Then we calculated the proportion of hypomethylated and hypermethylated genes for each patient. We categorized tumors for which greater than 1% of genes were hypermethylated as being “frequently hypermethylated” and tumors for which greater than 1% of genes were hypomethylated as being “frequently hypomethylated.” Frequent hypermethylation was more common (17.5% of tumors) than frequent hypomethylation (3.4%)(Figure 5). The maximum percentage of hypermethylated genes for any particular patient was 11.8%, compared to a maximum of 4.8% for hypomethylated genes.

**Figure 5:**
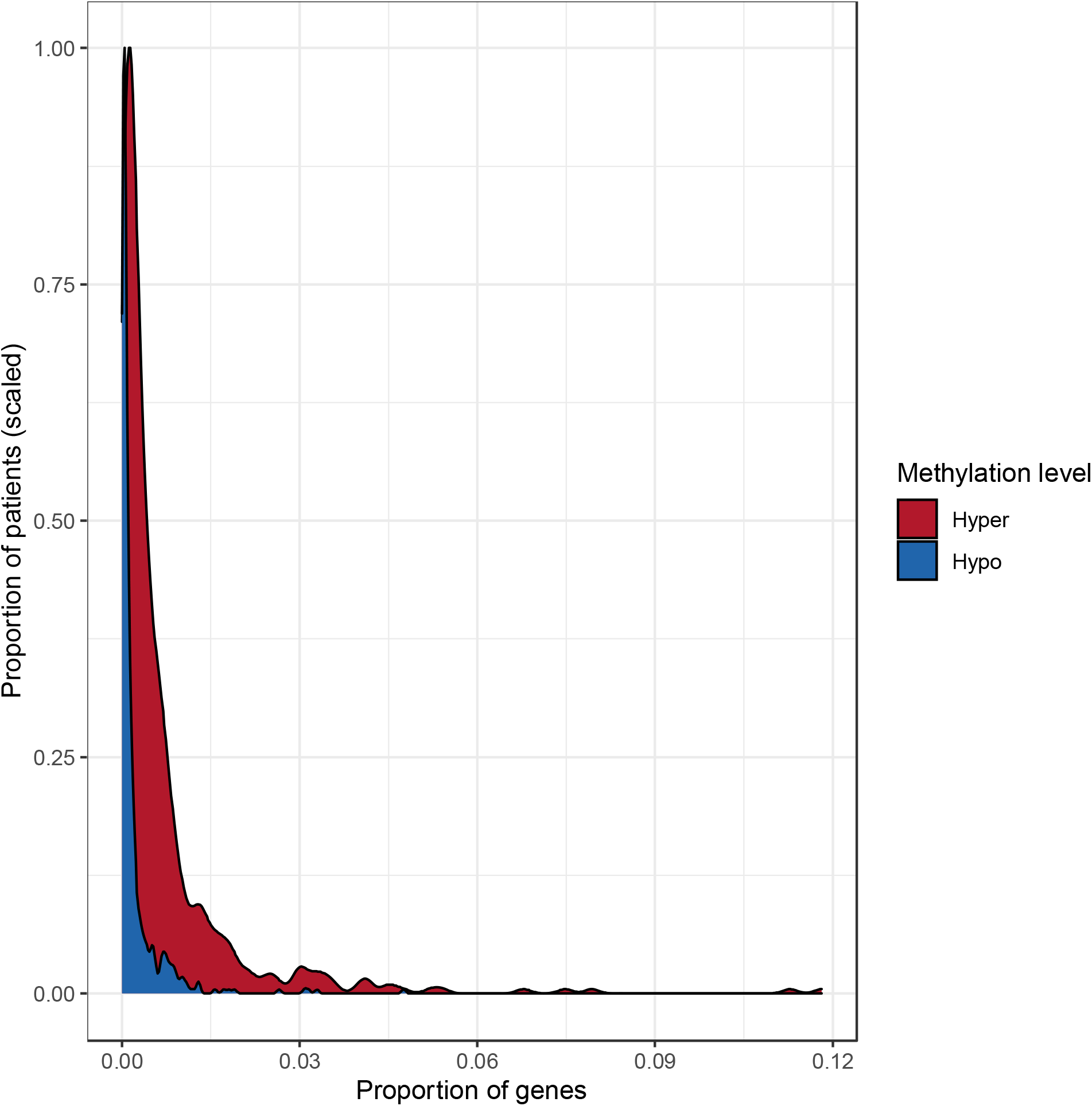
Distributions of the proportion of hypermethylated or hypomethylated genes in a given tumor. Using the normal data as a reference, we identified genes that were hypermethylated or hypomethylated in a given tumor. All five tumor types are represented. A relatively large number of hypermethylated genes in a given tumor was more common than a relatively large number of hypomethylated genes.

### Cancer mutation data analysis

Although they occur less frequently in pediatric tumors than in adult tumors, somatic mutations often contribute to pediatric tumorigenesis^4,5^. To evaluate the frequency of and interplay between somatic mutations and methylation events, we examined pediatric tumors for which both data types were available. Mutation data were available for somatic single nucleotide variants (SNVs), insertions/deletions (indels), and RNA fusions. We used the RNA fusion data as indicators of structural DNA variants. Copy-number data were available for only a small number of tumors, so we did not include this data type in the analysis. Methylation, SNV, indel, and RNA fusion data were available for Wilms tumors (n = 41), neuroblastomas (n = 65), and osteosarcomas (n = 66) but not for the other tumor types. For a given tumor, we considered genes with at least one SNV, indel, or RNA fusion event to be “mutated.” Across all tumor types, aberrant methylation—via either hypomethylation or hypermethylation of a given gene—occurred less frequently (in 1.1% of tumors) than mutations (2.8%); the largest disparity occurred for neuroblastomas (Table S3). Wilms tumors and osteosarcomas were aberrantly methylated nearly twice as often as neuroblastomas (1.4% vs. 0.8%; Table S3). In contrast, Wilms tumors and osteosarcomas were mutated less than half as often as neuroblastomas (1.7-1.8% vs. 4.6%).

Aberrant methylation levels and damaging mutations can have similar downstream effects in tumors^32^. After one of these alteration types has occurred in a given gene, it may be less likely for cells with a second alteration in the same gene to gain an additional selective advantage. Thus, having both a mutation and aberrant methylation in one gene may be less likely than expected by random chance. This mutual-exclusivity hypothesis has been examined extensively for pairs of genes in which somatic mutations might occur across diverse types of cancers^33–36^. It has also been investigated for DNA methylation events, though to a lesser extent^37^. Little is known about mutual exclusivity between methylation events and somatic mutations in pediatric tumors.

Treating each combination of tumor and gene as an independent observation, we examined whether DNA methylation events are mutually exclusive with somatic mutations. Mutations and aberrant methylation co-occurred rarely (0.03%) in the same gene and the same tumor (Table S3). Because both of these event types are rare, we reduced the data to the 962 genes for which at least 5 mutation events and at least 5 aberrant methylation events had occurred across the cancer types. We used a permutation test to evaluate whether mutations and aberrant methylation in the same gene and tumor co-occurred less frequently than would be expected by random chance. Across all tumors and the 962 genes, we observed 318 co-occurrences, whereas the average number of co-occurrences in the permuted data was 341.6. However, this difference was not statistically significant (p = 0.092).

### Oncogenes and tumor suppressor genes

Because oncogenes are typically expressed at relatively high levels and tumor suppressor genes are typically expressed at relatively low levels, we expected that oncogenes would have higher methylation levels than tumor suppressor genes. We identified 80 “tier 1” oncogenes and 141 “tier 1” tumor suppressor genes in the Cancer Genome Census^38^ and calculated the mean methylation value for each gene in the normal datasets and used a two-sample t-test to evaluate whether these mean values differed between the oncogenes and tumor suppressor genes. Under the hypothesis that oncogenes would be methylated at higher levels than tumor-suppressor genes, we used a one-sided test. The mean of the means for oncogenes was 0.02 higher than for tumor-suppressor genes; however, this difference was not statistically significant (p = 0.109). Many tumor-suppressor genes were highly methylated, and many oncogenes were methylated at low levels (Figure S1).

Next we evaluated the extent to which methylation levels differed for oncogenes and tumor-suppressor genes between tumor and normal conditions. First, we filtered the results from the two-sided Mann-Whitney U tests described above to include only oncogenes and tumor suppressor genes. Using FDR < 0.05 and an absolute mean difference > 0.02 as constraints, only 1 tumor suppressor gene was statistically significant (CTCF in Wilms tumors). No oncogenes reached statistical significance. We relaxed the threshold to FDR < 0.2 and removed the mean difference constraint. In this case, CTCF was significant for all cancer types except CCSKFigure S2. DNM2 (a tumor suppressor gene) was significant for OS, and GNA11 (an oncogene) was significant for NBLTable S4. Mean methylation values for none of the oncogenes or tumor suppressor genes differed across the tumor types (FDR < 0.2; Table S2).

Across the tumor types, mutation rates for oncogenes and tumor suppressor genes (0.031-0.089) were approximately twice the mutation rates for other genes (0.017-0.046) (Table 2), which aligns with prior evidence that mutations in these genes are associated with tumorigenesis. However, aberrant methylation rates for oncogenes and tumor suppressor genes (0.007-0.014) were approximately equal to methylation rates for other genes (0.008-0.014). In Wilms tumors and osteosarcomas, mutation rates for oncogenes and tumor suppressor genes were approximately three times higher than aberrant methylation rates, while mutation and methylation rates were approximately equal for other genes. In neuroblastomas, mutation rates for oncogenes and tumor suppressor genes were approximately twelve times higher than aberrant methylation rates but only six times higher for other genes.

**Table 2:** Aberrant methylation and mutation rates in oncogenes, tumor suppressor genes, and all other genes. For three pediatric tumor types, we identified aberrant methylation events (either hypomethylation or hypermethylation) that had occurred in a given tumor and gene. For the same tumor/gene combinations, we identified somatic single-nucleotide variants, indels, and structural variants that had occurred. These numbers indicate overall rates of aberrant methylation or mutation across all tumors of a given type. Methylation rates and mutation rates were typically similar across all three gene categories, but mutation rates for oncogenes and tumor suppressor genes were always higher than for other genes.

| **Tumor type** | **Aberration type** | **Oncogenes** | **TSG** | **Other** |
| --- | --- | --- | --- | --- |
| Wilms tumor | Mutation | 0.04 | 0.034 | 0.017 |
| Wilms tumor | Methylation | 0.012 | 0.012 | 0.014 |
| Neuroblastoma | Mutation | 0.083 | 0.089 | 0.046 |
| Neuroblastoma | Methylation | 0.007 | 0.007 | 0.008 |
| Osteosarcoma | Mutation | 0.031 | 0.034 | 0.018 |
| Osteosarcoma | Methylation | 0.014 | 0.011 | 0.014 |

We performed a modified version of the permutation-based, mutual-exclusivity analysis in which we searched for co-occurrences of either 1) a mutation in an oncogene and aberrant hypomethylation of the same gene or 2) a mutation in a tumor suppressor gene and hypermethylation of the same gene. Because few genes met these criteria, we performed this analysis with the oncogenes (n = 4) and tumor-suppressor genes (n = 29) that were mutated in at least 2 tumors and aberrantly methylated in at least 2 tumors. These event combinations were not mutually exclusive for oncogenes (p = 0.90) nor for tumor suppressors (p = 0.54).

## Discussion

Using publicly available data, we examined methylation patterns for five childhood cancers. We summarized baseline methylation levels for healthy children, identified deviations from these baseline levels in tumors, and evaluated mutual exclusivity of methylation events and somatic mutations. Subsequently, we evaluated these patterns for oncogenes and tumor-suppressor genes specifically. In the 309 healthy samples we studied, DNA methylation levels were highly consistent, suggesting that biological processes maintain this consistency in diverse tissue types and ancestral groups. In the 531 tumors we studied, hypermethylation or hypomethylation of promoter regions was a common feature, providing additional evidence that the tumor epigenome contributes to pediatric tumorigenesis. Hypermethylation was more common than hypomethylation. Hypermethylation of promoter regions has been associated with decreased gene expression^12^—tumors with frequent hypermethylation may result in broad silencing of proteins necessary for normal cellular function. Hypermethylation affecting multiple genes in the same tumor has been identified in adult thyroid neoplasms^39^ and colorectal tumors^40^ and has been associated with multidrug resistance in cell cultures^41^. However, global hypomethylation has garnered more attention than multigene hypermethylation^42^. Yet in the pediatric-tumor samples that we examined, widespread hypomethylation in a given tumor was uncommon. We also observed tumor subsets that exhibited divergent methylation—methylation levels for specific genes were consistently either high or low in many tumors of a given cancer type. While methylation dysregulation may be a common feature of pediatric tumors in general, specific mechanisms may drive these divergent changes in each tumor subset, and there may be overlap in these mechanisms across cancer types. Identifying such mechanisms may be useful for developing targeted treatments and informing patients.

Our mutual-exclusivity analysis did not provide evidence that somatic mutations and aberrant methylation are mutually exclusive with each other, whether for oncogenes and tumor-suppressor genes specifically or across all genes. However, we had access to a limited number of tumors (n = 172) for which both mutation data and methylation data were available; a larger-scale analysis is warranted. Furthermore, we evaluated single-nucleotide variants, indels, and structural variants because data were available for these mutation types. But large-scale amplifications and deletions occur regularly in tumors^43^, so including copy-number data in this type of analysis would also be useful in future research.

In normal cells, methylation levels did not differ significantly between known cancer genes (oncogenes and tumor suppressors) and other genes. In tumors, few oncogenes and tumor suppressors had high rates of aberrant methylation, suggesting that while aberrant methylation occurs in pediatric tumors, dysregulation likely does not occur disproportionately in these known gene categories. Global hypermethylation and/or hypomethylation patterns may be more important indicators of pediatric-tumor biology than local events. However, CTCF may be one exception. CTCF promoter methylation was significantly increased relative to normal conditions in every cancer type that we examined, except clear cell sarcoma of the kidney, which had a small sample size. CTCF codes for a zinc finger nuclease that is highly conserved in eukaryotes^44^. As a regulatory protein involved in DNA methylation^45^, improper function of this gene could lead to epigenetic abnormalities. Mutations in CTCF have previously been associated with several cancer types, including breast cancers, prostate cancers, and Wilms tumors^46,47^. Our findings across multiple pediatric cancer types suggest that CTCF hypermethylation—and consequent inhibition of gene expression— may influence pediatric tumorigenesis broadly and cause downstream effects.

In performing these analyses, we needed to choose arbitrary thresholds at times. For example, we specified three standard deviations above or below the mean in the normal data as a conservative threshold to indicate aberrant methylation. Using this threshold, we found that tumors were mutated more frequently than they were aberrantly methylated. However, if we had relaxed this threshold to two standard deviations, the average aberrant methylation rate would have been 0.045 rather than 0.012 (the average mutation rate was 0.027). Thus, our conclusions would have been moderately different. The use of arbitrary thresholds in research is common and cannot be completely avoided, especially when discipline-specific precedents have yet to be specified by the research community. In this scenario, we have introduced a new way of identifying aberrant methylation events. Future quantitative, experimental, and translational work will be necessary to improve our ability to determine when a particular gene in a given biological sample is methylated to an extent that alters that gene’s behavior and may in turn have clinical relevance.

A better understanding of DNA methylation’s role in pediatric tumors could shed light on mechanisms of tumorigenesis and eventually lead to insights about patient care and treatments. DNA methylation inhibitors have proven effective in some adult cancers, especially hematologic malignancies^20^ and may prove beneficial in pediatric cases. However, these therapies primarily target hypermethylation in a broad sense and thus may not be suitable for targeting specific genes^48,49^. Furthermore, as we have shown, hypermethylation events may be more common than hypomethylation events in pediatric tumors. Little is understood about how to reverse hypomethylation in vivo; however, global-hypomethylation patterns may be useful as biomarkers for therapies for targeting genes that have been activated as a result of hypomethylation.

## Methods

### Normal methylation datasets

We downloaded datasets containing DNA methylation levels for cohorts of normal patients as a way to establish a baseline against which tumor methylation could be compared for pediatric samples. We selected four datasets from Gene Expression Omnibus that used Illumina Infinium HumanMethylation450K arrays to profile normal cells in healthy fetuses and children aged 18 or under for which raw (.idat) files were available. Illumina Infinium HumanMethylation450K is a microarray platform that detects DNA methylation at over 450,000 locations in the human genome. Beta values from these arrays indicate the ratio of methylated signals to unmethylated signals^50^. Values closer to one indicate relatively high methylation levels; values closer to zero indicate relatively low methylation levels. Below we describe each dataset included in the study and provide accession identifiers from Gene Expression Omnibus.

GSE69502 originated from a study of Canadian second trimester fetuses^51^. Tissue was collected from chorionic villi, kidney, spinal cord, brain, and muscle. The original study analyzed DNA methylation differences in fetuses with spina bifida and anencephaly compared to normal fetuses. We included only data from the 65 normal fetuses.

GSE65163 originated from a study of nasal epithelial cells from African American children aged 10-12^52^. The original study analyzed DNA methylation differences in children with or without asthma. We included only data from the 36 children without asthma.

GSE109446 originated from a study of children aged 5-18 living in Cincinnati, Ohio (USA)^53^. Similar to GSE65163, nasal epithelial cells were used, and the goal of the original study was to understand methylation differences in children with or without asthma. We included only data from the 29 children without asthma.

GSE89278 originated from a study of Australian infant blood spot samples taken at birth^54^. The goal of the original study was to understand methylation differences in children with and without docosahexaenoic acid deficiency. We included only control data from 179 infants.

We chose these samples because they represented a variety of cell types and ancestral groups, which we hoped would make our findings more broadly generalizable than if our samples had been from a single, homogeneous population. Another factor in our decision was the age of the individuals. We searched for datasets representing relatively young patients to limit the possible confounding factor of methylation changes accumulating throughout life. In this process, we considered one additional normal dataset (GSE72556)^55^; however, we found that the beta levels from this dataset were consistently different from the other four normal datasets. This dataset originated from saliva samples, which often result in systematically different methylation levels than other types of samples^56–58^, perhaps due to external contaminants or different sample-collection procedures. As a result, we excluded this dataset from our analysis.

### Normal data processing

We processed the data using the minfi package (version 1.34.0) from R (version 4.0.2) and the Bioconductor suite^59,60^. We followed a standard workflow to process the methylation array files. The steps in this workflow included preprocessing, ratio conversion, and beta value calculations. We then summarized the beta values at the gene level. We mapped probes to genes based on an annotation file created by Price et al. (see http://www.ncbi.nlm.nih.gov/geo/query/acc.cgi?acc=GPL16304)61. Because we were interested in methylation changes in the promoter regions of genes, we included only probes\ within 300 base pairs of transcription start sites. We then calculated gene-level beta values for each patient as the mean beta value across all remaining probes in a given gene.

### Methylation data acquisition and processing

We obtained methylation data representing five tumor types from the Therapeutically Applicable Research to Generate Effective Treatments (TARGET) Data Matrix (retrieved August 24, 2020 from Website, https://ocg.cancer.gov/programs/target/data-matrix). We acquired data for 131 patients with Wilms tumor, 11 patients with clear cell sarcoma of the kidney, 68 patients with rhabdoid tumor, 235 patients with neuroblastoma, and 86 patients with osteosarcoma. We downloaded the data from TARGET using the rvest package (version 0.3.6)^62^. The data for Wilms tumors, clear cell sarcomas of the kidney, neuroblastomas, and osteosarcomas were generated using the Illumina Infinium HumanMethylation450K platform, and the data for rhabdoid tumors were generated using the Illumina Infinium MethylationEPIC platform. The HumanMethylation450K platform produces beta values for 22,579 genes, and EPIC produces values for 22,411 genes; we limited our analysis to the 22,253 genes that overlapped between the two platforms.

We followed the same workflow that we used to process the normal data. We calculated probe-level beta values using minfi and summarized values at the gene-level. We used principal component analysis to assess high-level patterns across the datasets and visualized the results using a scatter plot of the first two principal components. Methylation samples from each dataset generally clustered tightly with other samples from the same dataset, demonstrating that batch effects were presentFigure S3. To reduce the impact of these effects, we applied batch correction to a combined dataset that included all of the normal datasets. We used the dataset identifier as the batch variable. In addition, we logit transformed the data before performing the batch correction and inverse logit transformed the data after performing the batch correction to ensure the beta values stayed between 0 and 1. We performed these transformations using functions from the gtools package (version 3.8.2)^63^. Because the logit function cannot handle exact values of 0 or 1, we removed genes in which the beta values were exactly 0 or 1. We also removed genes in which beta values were NA. Filtering the data in this way removed 155 (0.7%) genes (out of 22,411 total genes). To perform the batch correction, we used the ComBat function from the sva package (version 3.36.0)^64^. After completing the batch correction, we performed another principal component analysis and visualized the results. The points no longer clustered by datasetFigure S3. We used these newly corrected values for all remaining analyses.

### Somatic-mutation data acquisition and processing

We obtained somatic-mutation data for TARGET patients via the Genomic Data Commons portal^65–67^. We first selected the Repository section. Under the Cases tab, we specified the TARGET program. We also specified Wilms tumor, neuroblastoma, and osteosarcoma as the tumor types of interest (data were unavailable for the other two tumor types). Under the Files tab, we selected “simple nucleotide variation” and “annotated somatic mutation” as the data category and type, and we indicated that we would use variants that had been called using Mutect2^68^. The data were stored in VCF format (version 4.2)^69^.

We wrote custom code to parse the mutation data for each patient. We included only mutations that had either 1) “HIGH” impact according to the variant annotations or 2) had “MODERATE” impact and were considered to be “deleterious” by SIFT^70^ or either “damaging” or “probably_damaging” according to Polyphen-2^71^.We considered using the specified minor-allele-frequency (MAF) values for filtering, but those values were unavailable for a significant portion of the variants, so we focused on functional annotations.

### RNA-fusion data acquisition and processing

We downloaded data for RNA fusions via the Genomic Data Commons portal. Under the Cases tab, we applied the same filters that we used for obtaining the somatic-mutation data. Under the Files tab, we selected “structural variation” and “Transcript Fusion” as the data category and type. We specified “STAR-Fusion”^72^ as the workflow type and “bedpe” as the data format. We wrote custom code to parse the RNA fusion data for each patient. The bedpe files described RNA transcripts from each patient that had genetic information originating from two separate genes. We considered an RNA fusion to have a functional impact on both genes affected by the fusion.

### Methylation and mutation data integration

Different naming conventions were used to identify neuroblastoma and osteosarcoma patients in the methylation data files versus the somatic-mutation and RNA fusion data files. We used a sample sheet provided by the Genomic Data Commons (gdc_sample_sheet.2021-03-11.tsv) and the sdrf metadata files for neuroblastoma and osteosarcoma to map the patient identifiers between these sources (see https://target-data.nci.nih.gov/Public/OS/methylation_array/METADATA/TARGET_OS_MethylationArray_20161103.sdrf.txt and https://target-data.nci.nih.gov/Public/NBL/methylation_array/METADATA/TARGET_NBL_MethylationArray_20160812.sdrf.txt). A few files did not map properly across naming conventions. One fusion data sample was mapped to two patients, so we excluded this sample. There were also two pairs of single nucleotide mutation samples that were mapped to single patients, so we excluded these samples.

### Inferring hypermethylation and hypomethylation states

To determine hypermethylation and hypomethylation status, we scaled the beta values for each gene to have a zero mean and a standard deviation of one across all normal samples. Then for each tumor sample, we compared the beta value for a given gene against the standardized values from the normal samples. If a tumor’s beta value was more than three standard deviations above the mean of the normal samples, we classified that tumor as being hypermethylated for that gene. If a tumor’s beta value was more than three standard deviations below the mean of the normal samples, we classified that tumor as being hypomethylated for that gene.

### Mutual exclusivity

We evaluated whether aberrant methylation—either hyper- or hypomethylation—was mutually exclusive with somatic-mutation events for a given gene. We calculated the total number of times a gene was both mutated and aberrantly methylated in the same tumor. Next we permuted methylation status for all tumors and kept mutation status constant. We repeated the permutation process 10,000 times to create an empirical null distribution to use as a baseline. For each permutation, we calculated the number of times a gene was both mutated and aberrantly methylated. We then compared the number of times that mutations co-occurred with aberrant methylation in the non-permuted data relative to the permuted data and calculated a p-value based on these numbers.

### Oncogenes vs tumor suppressor genes

We identified genes known to be oncogenes or tumor suppressor genes, based on information from The Cancer Gene Census^38^. We included only tier 1 genes classified as “oncogene” or “TSG” and ignored any for which it was ambiguous or unknown. To be classified as a tier 1 gene, there must be scientific evidence that a gene plays a role in cancer development and that mutations in the gene modify the activity of the associated protein. Several genes were listed in the Cancer Genome Census as both oncogenes and tumor suppressor genes; because of this ambiguity, we excluded these from our analysis.

In evaluating our hypothesis that oncogenes generally have decreased methylation levels in tumor samples relative to normal samples and that tumor suppressor genes generally have increased methylation levels in tumor samples relative to normal samples, we applied a two-sided, Mann-Whitney U test to each oncogene and tumor suppressor gene to identify genes that did (or did not) align with these expectations.

## Abbreviations

Abbreviation: Term
WT: Wilms tumor
CCSK: Clear cell sarcoma of the kidney
RT: Rhabdoid tumor
NBL: Neuroblastoma
OS: Osteosarcoma
TSG: Tumor suppressor gene

## Acknowledgements

We thank the research participants who donated samples for molecular profiling and the researchers who consented to have their data shared publicly. We thank the College of Life Sciences at Brigham Young University for providing funding to ACP through a College Undergraduate Research Award. The results published here are in whole or part based upon data generated by the Therapeutically Applicable Research to Generate Effective Treatments (TARGET) initiative, phs000218, managed by the NCI. The data used for this analysis are available from the National Cancer Institute Genomic Data Commons (https://gdc.cancer.gov). Information about TARGET can be found at http://ocg.cancer.gov/programs/target.

## Ethics approval and consent to participate

Brigham Young University’s Institutional Review Board approved this study under exemption status. This study uses data collected from public repositories only. We played no part in recruiting patients or obtaining consent.

## Competing Interests

The authors declare that they have no competing interests.

## Code and data availability

All of the code used to perform the analysis is publicly available on Open Science Framework so that others can verify and build upon our work (https://osf.io/79yfb/).

## Author Contributions

The contributions listed below correspond to the CRediT Taxonomy^73^.

ACP: Conceptualization, Formal Analysis, Funding Acquisition, Investigation, Methodology, Software, Visualization, Writing – Original Draft Preparation, Writing – Review & Editing

BIQ: Conceptualization, Formal Analysis, Software, Visualization, Writing – Original Draft Preparation, Writing – Review & Editing

SRP: Conceptualization, Funding Acquisition, Methodology, Project Administration, Resources, Supervision, Writing – Review & Editing

## Notes

https://osf.io/79yfb

### Competing Interest Statement

The authors have declared no competing interest.

