## Supplementary Material for "The DNA methylation landscape of five pediatric-tumor types"

### Supplementary Tables

**Table S1: Genes for which methylation levels differed between tumors and normal tissues.** We compared methylation levels at the gene level between each pediatric tumor type and the normal datasets. This table lists genes that resulted in a Bejamini-Hochberg False Discovery Rate of 0.05 or smaller and an absolute methylation change > 0.02.

| **Gene** | **p-value** | **FDR** | **Difference** | **Tumor type** |
| --- | --- | --- | --- | --- |
| KLHDC4 | 6.3e-25 | 1.4e-20 | -0.025 | Wilms tumor |
| CTCF | 3.8e-18 | 4.3e-14 | 0.031 | Wilms tumor |
| RNF146 | 9.0e-11 | 5.0e-07 | 0.045 | Wilms tumor |
| ZSCAN5A | 3.2e-09 | 1.4e-05 | 0.045 | Wilms tumor |
| LCA5L | 5.4e-08 | 1.7e-04 | -0.031 | Wilms tumor |
| FAM3B | 1.1e-07 | 2.4e-04 | -0.02 | Wilms tumor |
| GRID2 | 9.8e-08 | 2.4e-04 | 0.038 | Wilms tumor |
| MIR635 | 2.3e-07 | 4.7e-04 | -0.037 | Wilms tumor |
| MAEL | 5.7e-07 | 9.8e-04 | 0.029 | Wilms tumor |
| GLUL | 2.7e-06 | 3.8e-03 | 0.035 | Wilms tumor |
| TRH | 7.2e-06 | 9.0e-03 | -0.033 | Wilms tumor |
| PLXND1 | 9.7e-06 | 0.01 | 0.056 | Wilms tumor |
| FAM109A | 2.0e-05 | 0.02 | 0.034 | Wilms tumor |
| TPPP | 3.6e-05 | 0.03 | -0.032 | Wilms tumor |
| KLHL3 | 4.2e-05 | 0.03 | -0.034 | Wilms tumor |
| SDF4 | 1.7e-33 | 3.8e-29 | 0.065 | Rhabdoid tumor |
| UBE2J2 | 4.0e-28 | 4.5e-24 | -0.048 | Rhabdoid tumor |
| KCNQ1OT1 | 3.3e-23 | 2.5e-19 | -0.212 | Rhabdoid tumor |
| ETV6 | 6.5e-13 | 3.6e-09 | 0.045 | Rhabdoid tumor |
| LPP-AS2 | 1.9e-09 | 8.6e-06 | -0.022 | Rhabdoid tumor |
| PARS2 | 6.9e-07 | 2.5e-03 | -0.034 | Rhabdoid tumor |
| UBE2J2 | 2.1e-75 | 4.7e-71 | -0.079 | Neuroblastoma |
| KLHDC4 | 9.0e-46 | 1.0e-41 | -0.034 | Neuroblastoma |
| ZBED3 | 6.1e-36 | 4.5e-32 | 0.031 | Neuroblastoma |
| MIR635 | 4.6e-15 | 1.8e-11 | -0.033 | Neuroblastoma |
| KIAA1324 | 4.8e-10 | 1.5e-06 | -0.021 | Neuroblastoma |
| CDRT15P1 | 3.1e-09 | 6.9e-06 | -0.021 | Neuroblastoma |
| LOC642776 | 4.9e-09 | 9.9e-06 | 0.065 | Neuroblastoma |
| PXMP4 | 5.0e-08 | 7.4e-05 | 0.021 | Neuroblastoma |
| PLXNA3 | 5.9e-08 | 8.2e-05 | 0.082 | Neuroblastoma |
| GYS2 | 2.8e-07 | 3.5e-04 | 0.055 | Neuroblastoma |
| RAD54L2 | 8.6e-07 | 9.1e-04 | 0.032 | Neuroblastoma |
| SYCP2L | 6.8e-06 | 5.6e-03 | 0.022 | Neuroblastoma |
| PPP1R37 | 1.3e-05 | 9.2e-03 | -0.046 | Neuroblastoma |
| ZNF597 | 1.6e-05 | 0.01 | -0.034 | Neuroblastoma |
| C7orf66 | 1.7e-05 | 0.01 | -0.053 | Neuroblastoma |
| TDRD6 | 5.1e-05 | 0.03 | -0.024 | Neuroblastoma |
| TYR | 5.5e-05 | 0.03 | -0.048 | Neuroblastoma |
| PPP1R37 | 1.8e-12 | 3.9e-08 | -0.057 | Osteosarcoma |
| HDGFRP2 | 6.3e-09 | 7.0e-05 | 0.023 | Osteosarcoma |
| MIR635 | 5.8e-06 | 0.01 | -0.03 | Osteosarcoma |
| GDF9 | 9.9e-06 | 0.02 | -0.06 | Osteosarcoma |

**Table S2: Genes for which methylation levels differed significantly across pediatric tumor types.** We used one-way ANOVA tests to identify genes for which mean methylation values differed across the tumor types. This table shows genes that resulted in a Bejamini-Hochberg False Discovery Rate of 0.05 or smaller.

| **Gene** | **p-value** | **FDR** |
| --- | --- | --- |
| UBE2J2 | 7.3e-107 | 1.6e-102 |
| SDF4 | 1.7e-60 | 1.8e-56 |
| Metazoa_SRP | 1.4e-24 | 1.0e-20 |
| ZBED3 | 3.2e-23 | 1.8e-19 |
| TFAM | 2.8e-21 | 1.2e-17 |
| KLHDC4 | 9.0e-19 | 3.4e-15 |
| KCNQ1OT1 | 1.2e-17 | 3.8e-14 |
| LOC401093 | 3.3e-14 | 9.2e-11 |
| OTUD4 | 2.2e-10 | 5.4e-07 |
| KIAA1324 | 9.5e-09 | 2.1e-05 |
| LOC642776 | 2.0e-07 | 4.1e-04 |
| ZNF623 | 2.7e-07 | 5.0e-04 |
| FZD9 | 5.7e-07 | 9.7e-04 |
| LPP-AS2 | 6.1e-07 | 9.7e-04 |
| SBNO1 | 2.8e-06 | 4.2e-03 |
| TMSB4Y | 3.1e-06 | 4.3e-03 |
| VPS37A | 3.3e-06 | 4.3e-03 |
| NHSL2 | 7.9e-06 | 9.7e-03 |
| PLXNA3 | 1.0e-05 | 0.01 |

**Table S3: Frequency of aberrant methylation events, somatic mutations, and joint events affecting the same tumor and gene.** For three pediatric tumor types, we identified aberrant methylation events (either hypomethylation or hypermethylation) that had occurred in a given tumor and gene. For the same tumor/gene combinations, we identified somatic single-nucleotide variants, indels, and structural variants that had occurred. Next we quantified the number of times that both of these event types had affected the same tumor and gene. Numbers in the table represent all unique combinations of tumor and gene for which methylation data and mutation data were available.

| **Tumor type** | **Aberrantly methylated** | **Mutated** | **Mutated & aberrantly methylated** |
| --- | --- | --- | --- |
| Wilms tumor | 0.014 (12845/912373) | 0.017 (15182/912373) | 0.00024 (219/912373) |
| Neuroblastoma | 0.0075 (10856/1446445) | 0.046 (66312/1446445) | 0.00034 (486/1446445) |
| Osteosarcoma | 0.014 (20129/1468698) | 0.018 (26366/1468698) | 0.00028 (412/1468698) |
| Total | 0.011 (43830/3827516) | 0.028 (107860/3827516) | 0.00029 (1117/3827516) |

**Table S4: Oncogene and tumor suppressor genes for which methylation levels differed between tumors and normal tissues.** We compared methylation levels at the gene level between each pediatric tumor type and the normal datasets for oncogenes and tumor suppressor genes. This table lists genes that resulted in a Bejamini-Hochberg False Discovery Rate of 0.20 or smaller.

| **Gene** | **p-value** | **FDR** | **Difference** | **Tumor type** | **Gene category** |
| --- | --- | --- | --- | --- | --- |
| CTCF | 3.8e-18 | 4.3e-14 | 0.03 | Wilms tumor | TSG |
| CTCF | 4.9e-05 | 0.12 | 0.02 | Rhabdoid tumor | TSG |
| CTCF | 2.3e-08 | 3.9e-05 | 0.02 | Neuroblastoma | TSG |
| GNA11 | 3.7e-04 | 0.14 | 0.01 | Neuroblastoma | Oncogene |
| DNM2 | 8.2e-06 | 0.02 | 0.01 | Osteosarcoma | TSG |
| CTCF | 4.1e-05 | 0.07 | 0.02 | Osteosarcoma | TSG |

### Supplementary Figures

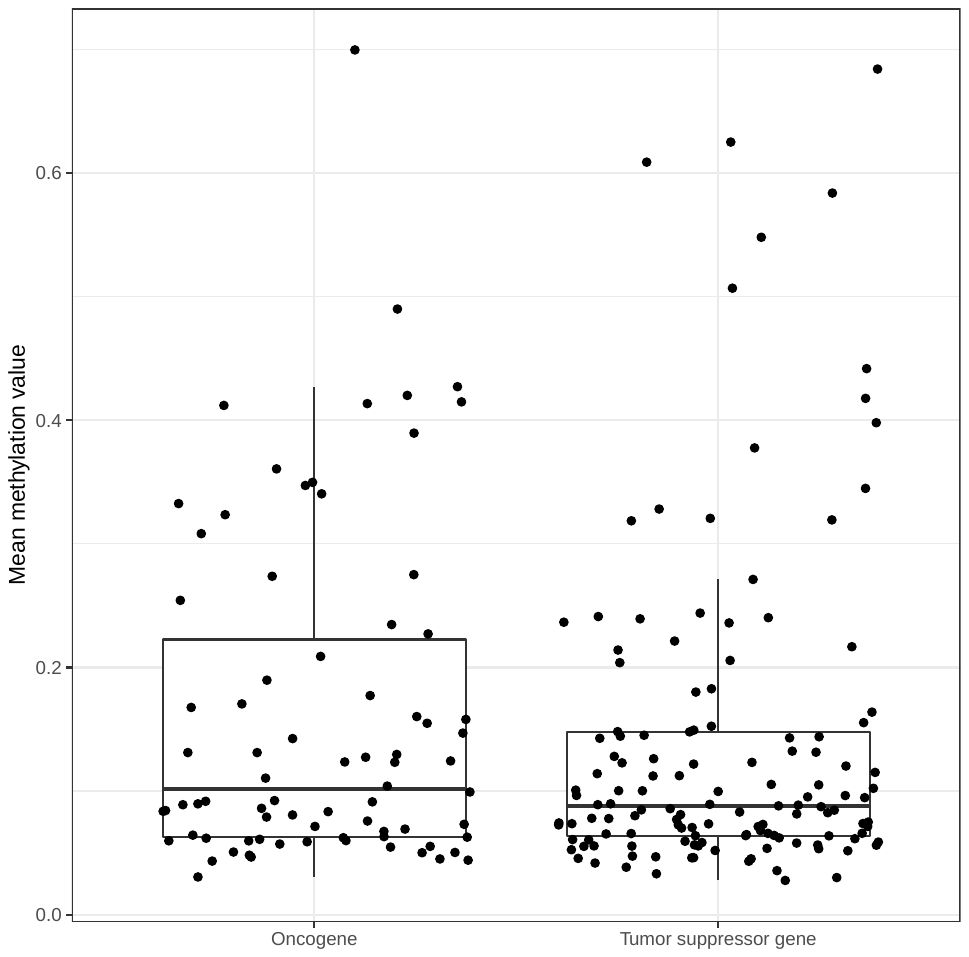

**Figure S1: Methylation levels for oncogenes compared to tumor suppressor genes.** We compared mean methylation levels under normal conditions for oncogenes and against levels for tumor suppressor genes. The difference between these two gene categories was not statistically significant.

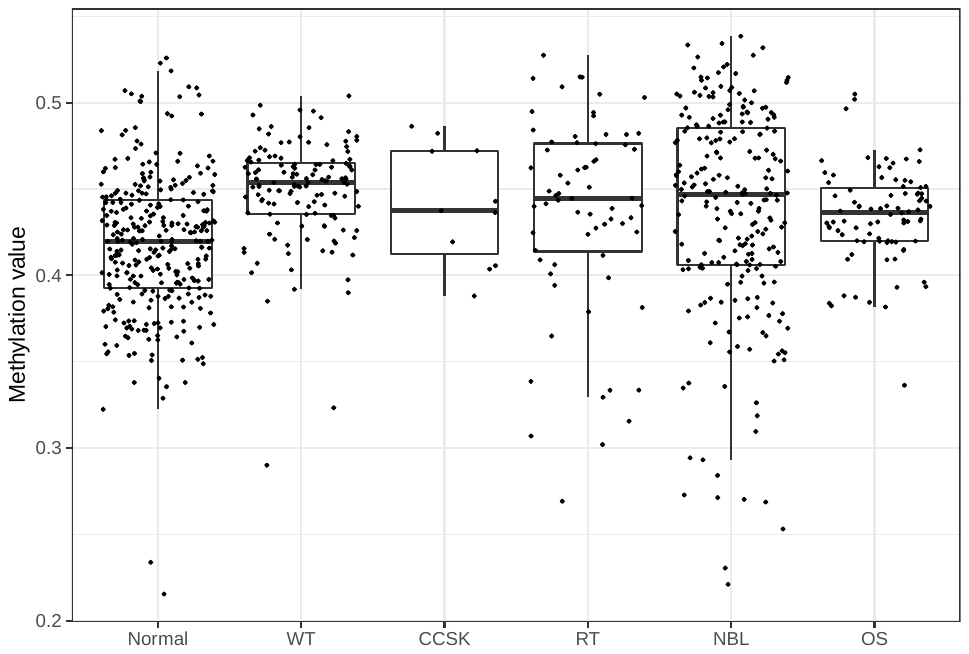

**Figure S2: Methylation levels for the CTCF gene under normal conditions and in tumors.** Methylation of CTCF was significantly higher than normal levels for all tumor types except clear cell sarcoma of the kidney. CTCF is a known tumor suppressor gene that regulates the epigenome.

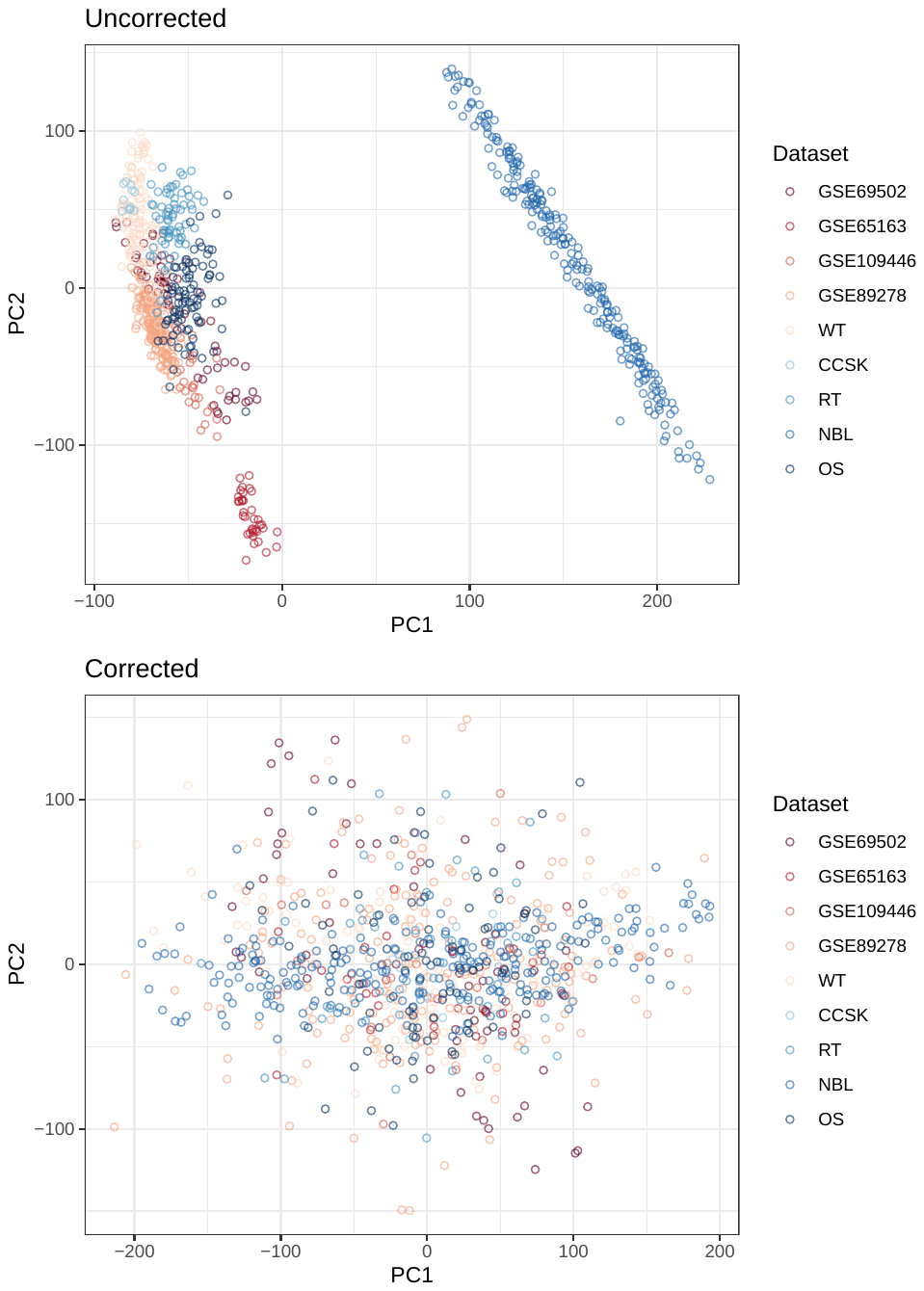

**Figure S3: Principal component analysis before and after batch correction.** A principal component analysis revealed that there were systematic differences in methylation levels across the datasets. Samples from each dataset typically clustered together. Some of this variance was expected, given that the samples originated from different tissue types. However, to support cross-tissue comparisons, including between normal samples and tumor samples, we used a linear model to adjust for these differences. Subsequently, a principal component analysis revealed that these systematic differences were no longer apparent in the data.

### Supplementary Additional Data Files

**Additional Data File S1: Methylation level / variance category for each gene in each normal dataset.**

**Additional Data File S2: Reactome pathway analysis results for Wilms tumors.**

**Additional Data File S3: Reactome pathway analysis results for rhabdoid tumors.**

**Additional Data File S4: Reactome pathway analysis results for osteosarcomas.**

**Additional Data File S5: Reactome pathway analysis results for neuroblastomas.**
